# Core-shell microparticle encapsulation for pH-responsive and targeted delivery of lactoferrin and ferrous sulfate

**DOI:** 10.64898/2025.12.10.693593

**Authors:** Claire E. Noack, Peilong Li, Waritsara Khongkomolsakul, Yunan Huang, Alireza Abbaspourrad

## Abstract

Microgel beads of amidated low methoxy pectin and bovine lactoferrin were formed by external gelation of a water in oil emulsion with ferrous sulfate. The stability of the lactoferrin to gastric digestion and proteolysis by pepsin was determined by gel electrophoresis. The microparticles were then dispersed in chitosan and the resulting mixture was spray dried to form a shell that is insoluble at neutral pH conditions. The iron content of the microparticles without chitosan was 34 mg g^-1^ and with chitosan was 27 mg g^-1^. The addition of chitosan lead to reduced iron release at pH 7 (30%) compared to 60% iron release without chitosan, but did not prevent iron from releasing in acidic conditions (pH 1). The core shell microparticle system shows promise as an iron fortificant in food applications.

## 1. Introduction

Iron is an essential micronutrient; it plays a vital role in oxygen transport, regulation of gene expression, and cell division (Abbaspour et al., 2014). Nearly 25% of the global population is anemic (Gardner et al., 2023). Anemia is highest among women of childbearing age and children (Sun et al., 2021). In these populations a chronic lack of iron can result in impaired growth and cognitive development, low birth weights, and neonatal fatality (Allali et al., 2017; Rahman et al., 2016).

The most common first-line treatment for iron deficiency anemia is oral supplementation (Auerbach & Adamson, 2016). Oral supplementation is affordable and simple to implement, but supplementation with high iron doses can lead to gastrointestinal distress (Tolkien et al., 2015). Oral supplementation with ferrous sulfate (FeSO_4_) is frequently used as it is inexpensive and has relatively high solubility (Allen, 2006). However, fortification of food with iron can lead to undesirable flavors and colors in food (Hurrell, 2002). Iron remains a challenging micronutrient for food fortification because less reactive forms of iron, such as elemental iron, have the disadvantage of low bioavailability due to poor solubility in the gastrointestinal tract, while high bioavailability iron, including ferrous sulfate, interact strongly with the food matrix (Allen, 2006).

Iron supplementation and absorption can be improved using iron encapsulation systems (Piskin et al., 2022). The encapsulation serves to decrease the undesirable effects of fortification of iron in food by reducing the ability of the iron to oxidize lipids, or interact with taste receptors (Kazemi-Taskooh & Varidi, 2021). Several approaches to iron encapsulation have been studied, including protein-iron gels relying on ionic gelation and spray drying with polysaccharides as the matrix material (Caetano-Silva et al., 2017; Muñoz-More et al., 2023; Pratap Singh et al., 2018). One such polysaccharide is pectin.

Pectin is extracted from apples and citrus fruits and often used in foods as a thickener or stabilizer (Lara-Espinoza et al., 2018). Pectin can form gels by cross-linking charged groups on the galacturonic acid monomers with iron ions through an egg-box model–a dimpled structure that surrounds the iron ions (Cao et al., 2020). Iron-pectin gels have been used to deliver iron (Ghibaudo et al., 2018), as well as protect and deliver probiotic bacteria to the small intestine (Ghibaudo et al., 2017). Pectin can be modified using ammonia under alkaline conditions to convert the methyl ester groups to amide groups (Reitsma et al., 1986). The amidation of pectin improves gelling ability by promoting hydrogen bonding and reduces sensitivity to ion concentration and pH (Said et al., 2023).

An alternate approach to improving Fe fortification is co-supplementation, where iron is dosed with a second compound in order to improve absorption and limit undesirable effects (Teucher et al., 2004). This includes prebiotic fibers, such as fructo- and galacto-oligosaccharides, which serve to decrease gut pH, improving the solubility of iron in the intestinal system (Husmann et al., 2022). Ascorbic acid is commonly supplemented with ferrous iron to serve as an antioxidant, preventing the formation of less soluble and less bioavailable ferric ions from ferrous ions (Li et al., 2020). Iron absorption can also be improved through co-supplementation with bovine lactoferrin (LF) (Mikulic et al., 2020).

LF is an iron binding glycoprotein in the transferrin family produced by mammals, including humans and cows, with high sequence similarity (Baker & Baker, 2004). LF is a bioactive protein, exhibiting antibacterial, antiviral, and anticancer activity (Kowalczyk et al., 2022). LF has been shown to interact with protein receptors in the duodenum, increasing iron absorption beyond supplementation with ferrous sulfate (Suzuki et al., 2005). Unfortunately, LF is susceptible to gastric proteolysis by pepsin in the stomach (Troost et al., 2001; Wang et al., 2017). While there are bioactive peptides produced during the hydrolysis of LF, notably lactoferricin, many of the functions of LF are dependent on an intact structure (Furlund et al., 2013; Wang et al., 2019). Therefore it is important to deliver the intact lactoferrin to the duodenum..

Bioactive delivery can be targeted to specific areas of the gastrointestinal tract through the choice of wall material. For example, enteric coatings are insoluble in acidic gastric conditions, but release their payload in neutral to alkaline conditions (Maderuelo et al., 2019). Examples of enteric coatings used in micronutrient delivery include the use of the polymer Eudragit S100 where the release of vitamins A, B2, C, D, and E was successfully targeted to the colon (Pham et al., 2021). Reverse enteric coatings, as their name implies, are insoluble at neutral pH conditions but dissolve readily in acid. Reverse enteric coatings have been used for targeted delivery of iron and iodine in double-fortified salt (Dueik & Diosady, 2017). Reverse enteric coatings can be formed with synthetic, acid-soluble polymers such as Eudragit EPO or natural polymers such as chitosan (Pratap Singh et al., 2018). For iron delivery, reverse enteric coatings are an improvement over enteric coated, or slow-release iron supplements because iron absorption takes place primarily in the duodenum, at the start of the small intestine while enteric coatings would release iron through the small intestine and colon (Stoffel et al., 2020).

Chitosan is formed by the deacylation of chitin, a polymer extracted from crustacean shells (Aranaz et al., 2021). Chitosan can also be produced by bacteria, removing concerns about allergens and kosher food safety (Abdel-Gawad et al., 2017). Chitosan is available in a variety of molecular weights, below 100 kDa is considered low molecular weight, 100-1000 kDa medium, and above 1000 kDa is considered high (Gonçalves et al., 2021). It is soluble below pH 4, but shows very limited solubility at pH 7 (Sogias et al., 2010).

Chitosan can be used as a matrix wall material to protect the bioactives of a core-shell microparticle. Core-shell microparticles can be produced by spray drying (Galogahi et al., 2020). Spray drying is a common technique in microencapsulation due to its high throughput, limited damage of heat labile components, and easy scale up (Gharsallaoui et al., 2007).

Here, we present an iron and LF co-supplementation vehicle formed by gelation of the dispersed phase of a water in oil emulsion. The microparticles were spray dried with or without chitosan. The stability of LF against gastric digestion was measured, as well as the release of iron in acidic or neutral pH conditions.

## 2. Materials and Methods

### 2.1 Materials

Lactoferrin (Bioferrin 2000; Iron >15 mg/100g) was provided by Glanbia Nationals, Inc, (Fitchburg, WI). Amidated low-methoxyl pectin (aLMP) was provided by CP Kelco (DE 27%, DA 20%, USA). Coconut medium-chain triglyceride (MCT) oil was purchased from Cocojojo Organic (FCC, Irvine, CA, USA). Ferrous sulfate was purchased from MP Biomedicals (>99%, Santa Ana, CA, USA). Soy lecithin was purchased from Tokyo Chemical Industry (ACS, Tokyo, Japan). Nile Red was purchased from Chem-Impex (ACS, Wood Dale, IL, USA). Fluorescein isothiocyanate isomer I (FITC) was purchased from Sigma-Aldrich (>90%, St. Louis, MO, USA)

### 2.2 Zeta potential measurement

The zeta potential was measured using a Malvern Zetasizer Nano ZS (Malvern, Germany). aLMP (2 mg mL^−1^) was adjusted to the target pH in 1.0 unit increments with 1 M HCl or 1 M NaOH. The microparticles were measured at 1 mg mL^−1^ in MilliQ water. Each measurement was taken in triplicate with >10 runs per sample in Smoulchwski mode.

### 2.3 Iron encapsulation efficiency

The interaction between aLMP and Fe was measured by the encapsulation efficiency of 2 mM FeSO_4_ by 2 mg mL^−1^ aLMP. The biopolymer and Fe(II) ions were mixed for 30 min, then the solution was centrifuged for 10 min at 10000 g. The iron concentration in the supernatant was determined colorimetrically by the ferrozine method (Carpenter & Ward, 2017). A standard curve of 0.02 mM to 0.10 mM FeSO_4_ was used to calculate the total Fe concentration in the samples. The encapsulation efficiency was calculated by Eqn. 1

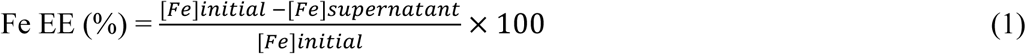

### 2.4 aLMP-Fe gel formation

LF (25 mg mL^−1^) and aLMP (25 mg mL^−1^) were hydrated for 2 h. The solutions were adjusted to pH 6 using 1 M NaOH. The aLMP-Fe gel was formed by mixing 10 mL of aLMP with the appropriate volume of 1 M FeSO_4_ to result in 10 mM, 50 mM, or 100 mM FeSO_4_ in the final solution. The same process was followed for the LF-aLMP-Fe gel, with the biopolymers mixed in a 1:1 volume ratio prior to the addition of iron. The gels were allowed to cure for 24 h before observation.

### 2.5 Preparation of microparticles

LF and aLMP (500 mg of each) were separately dissolved in 20 mL MilliQ water for 2 h at room temperature. The solutions were adjusted to pH 6 using 1 M NaOH, then mixed. This aqueous phase (40 mL) was emulsified with 160 mL of coconut MCT oil containing 1.6 g of soy lecithin. The mixture was homogenized with a high speed homogenizer (T25 digital Ultra-Turrax, IKA, Germany) for 2 min at 6000 rpm. The resulting water in oil emulsion was transferred to a mechanical stirrer and 2 mL of 1 M FeSO_4_ solution was added dropwise at 1 mL min^−1^. The emulsion was stirred for 30 min to allow the dispersed phase to gel. To remove the oil phase, the emulsion was centrifuged twice at 2000*g* for 5 min. The pellets were then frozen at – 20 °C overnight, then freeze dried for 48 h using a Freezone 1 lyophilizer (Labcono, Kansas, MO, USA) with a moisture collector temperature of −50 °C and a vacuum pressure of −0.049 mBar

### 2.7 Particle size measurement

The particle size and distribution of the emulsion droplets was determined using a Malvern Zetasizer Nano ZS (Malvern, Germany). The water-in-oil (w/o) emulsion was diluted 100x with more MCT oil to aid in accuracy of measurement. The particle size was measured before the addition of iron and after allowing the water droplets to gel. The parameters were set with a refractive index of 1.45 for MCT oil and a backscattering angle of 173°.

### 2.8 Confocal laser scanning microscopy

The continuous and dispersed phases of the emulsion were visualized using a confocal laser scanning microscope (i880, Carl Zeiss, Göttingen, Germany). 50 µg mL^−1^ Nile red was added to the MCT oil prior to emulsification to act as a liposoluble fluorescent probe. The LF solution was labeled with fluorescein isothiocyanate (FITC) prior to emulsification to allow visualization of the protein within the emulsion. LF was labeled with FITC by dissolving FITC at 1 mg mL^−1^ in DMSO and adding it dropwise to a solution of 5 mg mL^−1^ of LF in 0.1 M sodium bicarbonate buffer at pH 9. The FTIC and LF were allowed to interact at 4 °C overnight, and the residual free FITC was removed by dialysis against MilliQ water for 24 h, changing the water after 0.5, 2, 6, and 24 h. The FITC-labeled LF was freeze dried prior to use. The emulsion was examined with an excitation/emission wavelength of 561/654 nm for Nile red and 488/522 nm for FITC, with and without brightfield.

### 2.9 Spray Drying

After oil removal, the collected aqueous phase pellet was dispersed in 100 mL MilliQ water containing 30 mg mL^−1^ of maltodextrin. For chitosan-containing microparticles, 15 mg mL^−1^ of chitosan was dispersed with the maltodextrin solution. The microparticles were then spray dried using a bench mounted spray dryer (Armfield SD-basic, FT30MkIII, Hampshire, England) with an inlet temperature of 180 °C, an outlet temperature 70 °C, and a feed rate of 10 mL min^−1^.

### 2.10 Lactoferrin Loading

The concentration of LF in the microparticles was quantitatively determined by the Pierce Rapid-Gold bicinchoninic acid assay (BCA) using bovine serum albumin (BSA) as the standard. 10 mg mL^−1^ of LF-aLMP-Fe or CS-LF-aLMP-Fe microparticles were dispersed in 1 M HCl to improve solubility. The LF loading was calculated by Eqn. 2

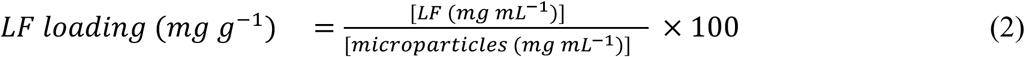

### 2.11 Scanning electron microscopy

The microstructure of the spray dried microparticles was analyzed using a Zeiss Gemini scanning electron microscope (Zeiss, DE). The samples were placed on a carbon taped stub and coated using a Denton Desk V sputter coater (Denton, NJ, USA) with carbon for 30 s. The images were taken using a 20 µm aperture and an accelerating voltage of 3.0 kV on an HE-SE2 (backscattered) detector.

### 2.12 In Vitro Digestion Model

The stability of the LF against gastric digestion was tested using the INFOGEST 2.0 method (Brodkorb et al., 2019). Briefly, 80 mg of LF, or the equivalent weight of microparticles that contains 80 mg of LF was dispersed in 2 mL of simulated salivary fluid (SSF). The sample was heated at 37 °C with agitation for 2 min. Following, 4 mL of simulated gastric fluid (SGF) with pepsin was added and the solution was adjusted to pH 3 with 1 M HCl. The mixture was kept at 37 °C with agitation for 120 min. Aliquots (1 mL) were removed at 10 min, 30 min, and 120 min and adjusted to pH 7 with 1 M NaOH to stop enzyme activity. To the remaining 5 mL of SGF, 5 mL of simulated intestinal fluid with pancreatin was added and the pH was adjusted to 7 with 1 M NaOH. Aliquots (1 mL) were removed at 10 and 120 min and heated to 95 °C for 5 min to terminate enzyme activity.

The remaining LF content after digestion was confirmed qualitatively by SDS Page. The protein samples were diluted to approximately 2 mg mL^−1^ and mixed in a 1:1 ratio with 2x Laemmli sample buffer and heated at 95 °C for 5 min. The sample (10 µL) was added to each well. The SDS Page was performed with a BIORAD Rapid Cast Kit 12% polyacrylamide gel, at a voltage of 80 kW through the stacking gel and 110 kW through the resolving gel in a running buffer of 25 mM Tris base, 0.19 M glycine, and 3.46 mM SDS. The samples were dyed with 150 mg mL^−1^ of Comassie Brilliant R-250 in a solution consisting of 500 mL L^−1^ methanol, and 100 mL L^−1^ acetic acid for 1 h, then destained in a buffer containing 200 mL L^−1^ methanol and 100 mL L^−1^ acetic acid for at least 12 h.

### 2.13 Iron bioavailability and release

The iron release behavior of the spray dried microparticles was analyzed according to the method previously described (Pratap Singh et al., 2018). Briefly, microparticles containing 2 mg Fe were added to 5 mL of 0.1 N HCl or 10 mM pH 7 potassium phosphate buffer. The samples were agitated gently in a 37 °C shaking water bath. 100 µL aliquots were removed at 15, 30, 45, 60, 90, and 120 min. The concentration of free iron measured in the aliquot after centrifugation at 10,000*g* for 5 min was determined colorimetrically by the ferrozine method. The percent of iron released at 30 min in 0.1 M HCl is taken to be the in vitro bioavailability of the iron (Swain et al., 2003). Iron release was calculated using Eq. 3:

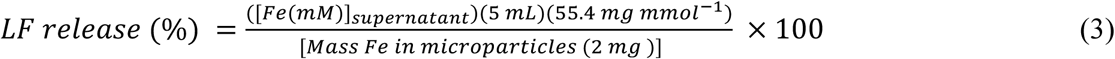

## 3. Results and Discussion

### 3.1 Iron-pectin interaction and gel formation

The interaction between iron ions and pectin was used to trap bioactives in the resulting gel matrix. The iron-induced gelation of aLMP is pH dependent, with an Fe encapsulation efficiency (EE) of nearly 40% at pH 7, but only 20% at pH 2 (**Fig 1A**). The observed pH dependence of the EE is likely due to the increased negative charge on the aLMP as the pH approaches neutral (**Fig 1B**). The charge density of aLMP is a result of the deprotonation of the carboxyl groups on the galacturonic acid residues on the polymer chain (Gallery et al., 2024). The pKa of the carboxyl groups in pectin is 3.5, above which the negative charge of the carboxylate allows it to form an egg-box like structure with the ferrous ions. The small observed encapsulation of iron at pH 2, when the zeta potential of pectin is positive, can be explained by aLMP’s ability to form gels in acidic solutions with or without cations present, however without the additional negative charges to attract additional ferrous ions encapsulation is poor (Capel et al., 2006).

**Fig 1.**
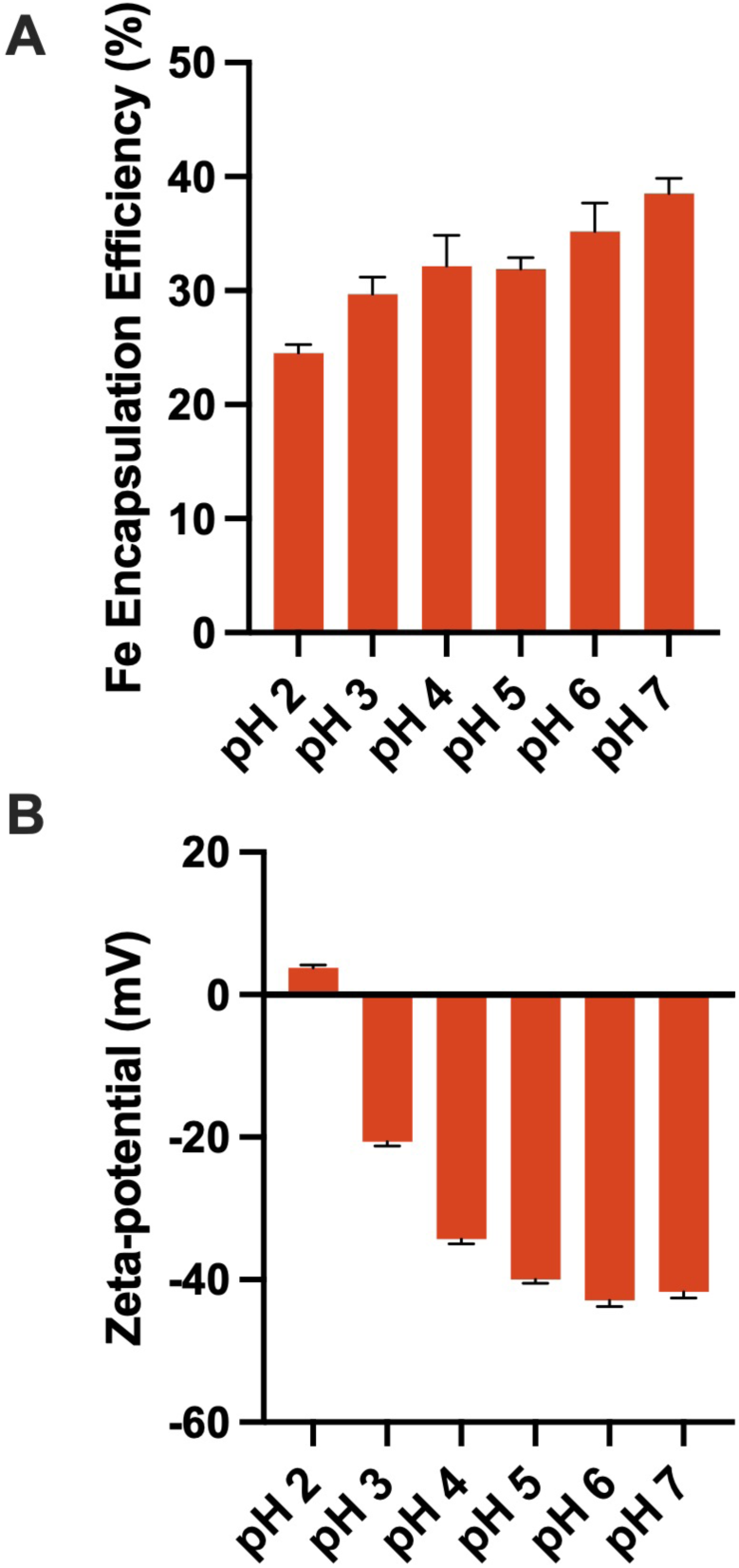
Fe(II) encapsulation of 10 mM FeSO_4_ with 2 mg mL^−1^ aLMP in MilliQ water at pH 2-7 (A). Zeta potential of 2 mg mL^−1^ aLMP in MilliQ water at pH 2-7 (B).

The strength of the iron-pectin gel and the ability to trap LF is dependent on the iron concentration in the gel. aLMP can form gels in a wide range of cation concentrations (Said et al., 2023). To determine the appropriate range for gelation, iron sulfate was added to a mixture of 1:1 LF to aLMP to achieve Fe(II) concentrations of 10 mM, 50 mM, and 100 mM Fe(II) (**Fig. S1**). At 10 mM Fe(II), the sample did not form a complete gel; free water surrounded the gelled center. Both 50 mM and 100 mM were sufficiently high to form a gel and traptrap the water and LF. However, the 100 mM gel was brittle leading to the conclusion that 100 mM Fe resulted in over-crosslinking of the gel, decreasing extensibility and increasing stiffness (Ngouémazong et al., 2012). Because the upper recommended limit for iron intake is 40 mg Fe per day (EFSA Panel on Nutrition, Novel Foods and Food Allergens (NDA) et al., 2024) (EFSA Panel on Nutrition, Novel Foods and Food Allergens (NDA) et al., 2024), 50 mM Fe as the optimum concentration for further experiments.

### 3.2 Water in oil emulsion gelation

The LF-aLMP-Fe microparticles were produced by external gelation of the dispersed phase of a water-in-oil (W/O) emulsion (**Fig. 2A**). Preliminary experiments were performed to determine the optimum water:oil ratio, emulsifier, and gelation time. The water to oil ratio was adjusted to result in an emulsion with a liquid, instead of semi-solid texture. Soy lecithin was chosen because it resulted in the formation of an O/W emulsion, rather than W/O. Allowing the dispersed phase to gel longer than 30 min did not promote further gelation. The current formulation resulted in a gelled aqueous phase with clear removal from the oil phase after centrifugation. After oil removal, the pellet was able to be re-dispersed in MilliQ water for further characterization or spray drying

**Fig 2.**
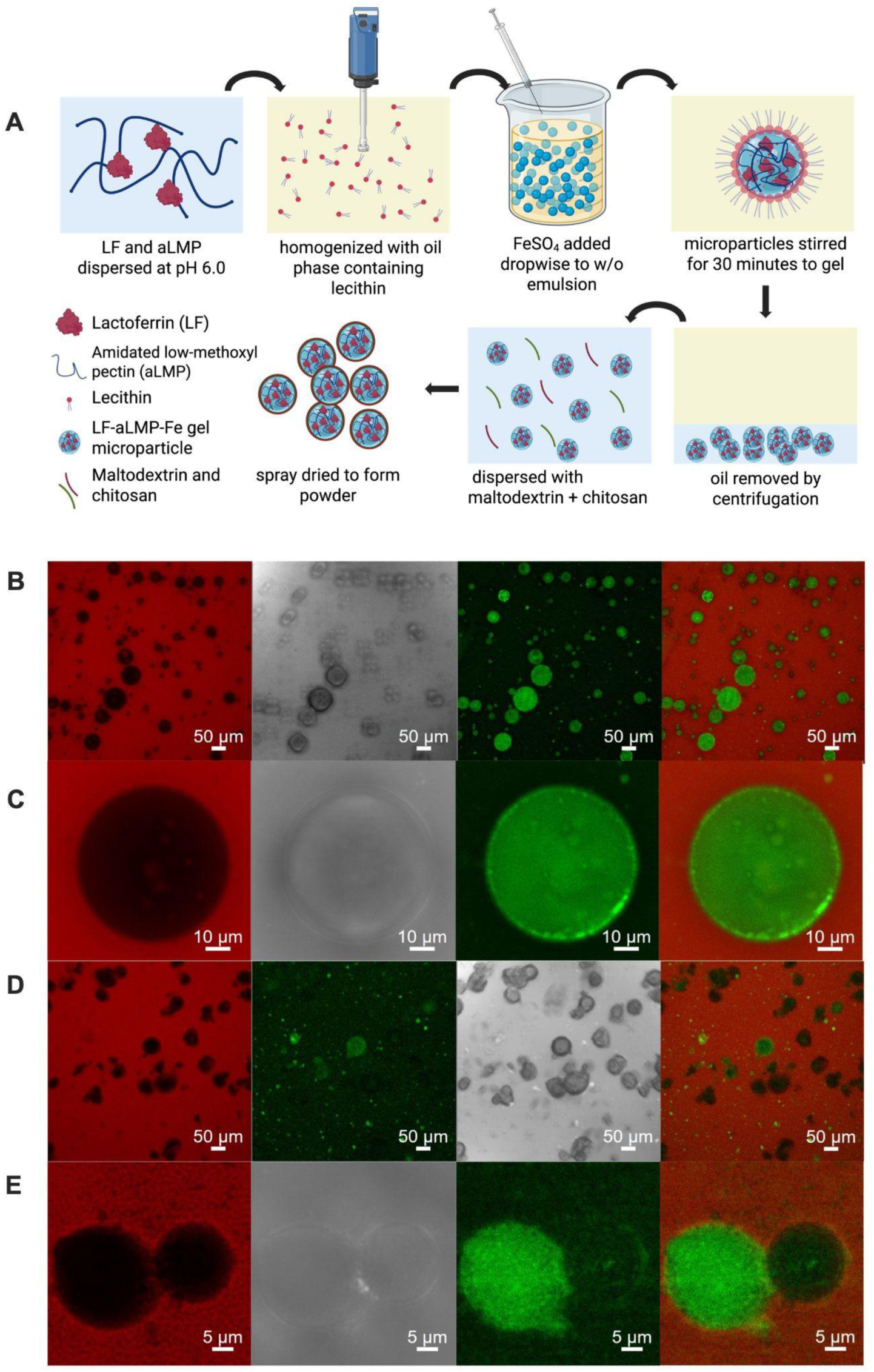
Emulsion gelation process of LF-aLMP-Fe microparticles (A). CLSM images of the initial emulsion (B,C) and iron gelled emulsion droplets (D,E). Photos were captured in different emission channels for Nile red (red) and FITC (green).

Confocal laser scanning microscopy was used to identify the continuous and dispersed phases of the emulsion (**Fig. 2B**). The initial emulsion can be identified as a W/O emulsion with a non-uniform dispersed phase droplet size. Additionally, inclusions of oil can be seen in the aqueous phase, showing an O/W/O double emulsion. This is likely due to the emulsifying activity of LF, which has been shown to stabilize O/W emulsions as successfully as Tween 20 (Teo et al., 2016).

Closer inspection of one aqueous droplet shows the LF fluorescence is higher on the interface of the system (**Fig 2C)**. This is again owing to the surface activity of LF, where hydrophobic patches on the protein will be adsorbed onto the oil-water interface to decrease surface tension. A Z-stack of the fluorescence signal of the droplet shows that while the LF is concentrated on some parts of the interface, it is still distributed throughout the sphere (**Fig S2)**.

After the addition of FeSO_4_ the aqueous phase is no longer spherical, indicating that a gel has been formed (**Fig 2D**). In emulsified systems the dispersed phase normally forms a sphere to minimize interfacial tension. The non-spherical appearance of the aqueous phase indicates that the gel is holding the shape of the water droplet. The LF green fluorescence signal is lower in the gelled emulsion compared to the initial emulsion. FITC fluorescence is strongly affected by the presence of iron ions, and the presence of Fe(III) will quench the fluorescence signal (Zhou et al., 2014). Some of the water droplets show a higher fluorescence signal than others (**Fig 2E)**. The distribution of the fluorescence signals within the water droplets was not homogeneous, we attribute this to potential factors: an uneven distribution of iron ions between the various water droplets, or the iron solution forming droplets in the aqueous phase with itself rather than cross-linking and gelling the droplets containing LF and aLMP.

### 3.3 Microparticle characterization

The particle size distribution and surface charge of the biopolymers and microparticles were measured to determine their colloidal stability and suitability for spray drying (**Table 1**). aLMP is an anionic polymer at pH values above the pKa of its carboxyl groups, around 3.5 with an overall surface charge of -49 mV at pH 6 (Cao et al., 2020), while LF has a high isoelectric point (pI) of pH 8, and as such carries a positive overall surface charge of 14.3 mV at pH 6 (Baker & Baker, 2004). The charge of the LF-aLMP-Fe microparticles at pH 6 was -10.2 mV, less negative than the charge of aLMP alone due to the addition of the positive Fe(II) ions and positive LF.

**Table 1.** Zeta potential and particle size of biopolymers and gel microparticles.

| Sample | Zeta potential (mV) | Z-average (nm) | PDI |
| --- | --- | --- | --- |
| LF | $14.3 \pm 0.5$ | 122 | 0.25 |
| aLMP | $-49.8 \pm 1.3$ | 1933 | 0.32 |
| CS | $52.6 \pm 0.6$ | 1467 | 0.61 |
| LF-aLMP-Fe | $-10.2 \pm 0.1$ | 3207 | 0.40 |
| CS-LF-aLMP-Fe | $31.0 \pm 0.5$ | 1121 | 0.78 |

Chitosan (CS) is unique among nature-derived biopolymers because it carries a positive charge (Jiménez-Gómez & Cecilia, 2020). CS can interact favorably with the negative surface of the LF-aLMP-Fe microparticles due to the electrostatic attraction between the opposite charges (Niu et al., 2019). The resulting surface charge of the microparticles with CS added is highly positive, at 31 mV.

The particle size of the biopolymers and microparticles was measured using dynamic light scattering to determine the suitability of the dispersions for spray drying. The size of dispersed particles needs to be much smaller than the spray dryer orifice to prevent clogging (Malamatari et al., 2020). The Z-average particle size is most accurate with monodisperse samples with a polydispersity index (PDI) below 0.3. The PDI of aLMP and CS shows that the biopolymers were non-uniform but large in size. The average particle size of the LF-aLMP-Fe microparticle was larger than the CS-LF-aLMP microparticle. This is likely due to aggregation of several microparticles together. The zeta potential of a particle in solution plays an important role in the stability of a dispersion, with zeta potential values higher than +30 mV or lower than - 30 mV having a high degree of stability, but the range of ± 10-20 mV as relatively stable (Cano-Sarmiento et al., 2018).

### 3.4 In vitro digestion stability

The INFOGEST protocol is a static digestion model that can be used to show the stability or breakdown of proteins in the gastrointestinal tract (Brodkorb et al., 2019). The first phase of digestion, the salivary phase, contains amylase enzymes at a pH of 7. LF is not affected by amylase activity, and so was not measured at this time point (van der Maarel et al., 2002). Samples were treated with pepsin at pH 3, and aliquots were taken after 10, 30, and 120 min, as well as after 10 and 120 minutes in the simulated intestinal phase at pH 7 with pancreatin. The digestion of LF was followed by SDS-PAGE (**Fig 3**). The non-encapsulated LF was extensively degraded even after 10 min in the gastric phase and no intact LF was observed in the intestinal phase. Trapping the LF within the aLMP-Fe bead showed protection through the gastric phase, with some LF remaining after 10 min in the intestinal phase.

**Fig 3.**
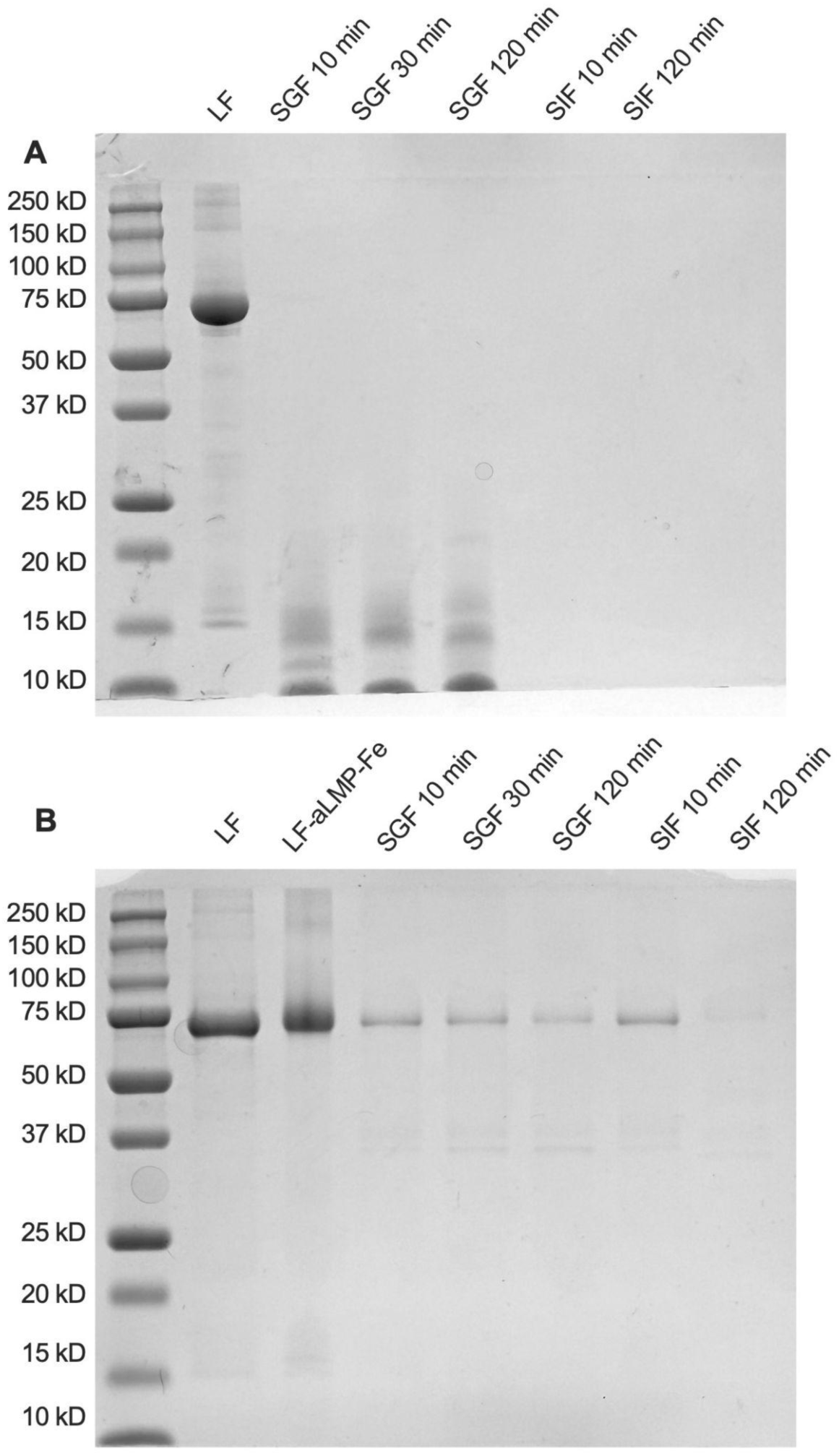
SDS-PAGE images of INFOGEST simulated digestion fluids at SGF 10 min, SGF 30 min, SGF 120 min, SIF 10 min and SIF 120 min

Preservation of the LF structure through the gastric phase is important because the site of activity of iron absorption is in the duodenum, at the start of the small intestine (Gulec et al., 2014). LF receptors have been proposed that help with iron absorption in the proximal small intestine (Suzuki et al., 2005). The SDS-PAGE results indicate that intact LF can be delivered to the small intestine through encapsulation.

### 3.5 Spray dried microparticle characterization

Spray drying dispersions into powders involves high temperatures and rapid water removal. The short time of exposure to heat is preferred in industry to preserve structures of heat-labile proteins such as LF (da Silva Júnior et al., 2023). The LF-aLMP-Fe and CS-LF-aLMP-Fe microparticles were re-dispersed in aqueous solutions of 30 mg mL^−1^ maltodextrin as the spray drying carrier. Maltodextrin is commonly used as a bulking agent in spray drying to produce a flowable, less sticky powder (Nadali et al., 2022). The spray dried powders were examined using a scanning electron microscope (SEM) (**Fig. 4)**. The spray dried powder of LF alone shows buckling of the powder, which can happen as a crust forms at the outset of drying as the moisture in the interior of the droplet escapes (Rogers et al., 2012). LF-aLMP-Fe microparticles show less buckling, and more of a spongy texture. The CS coated microparticles showed the appearance of some “puffed” particles, potentially as a result of higher solids in the feed solution resulting in better particle integrity.

**Fig 4.**
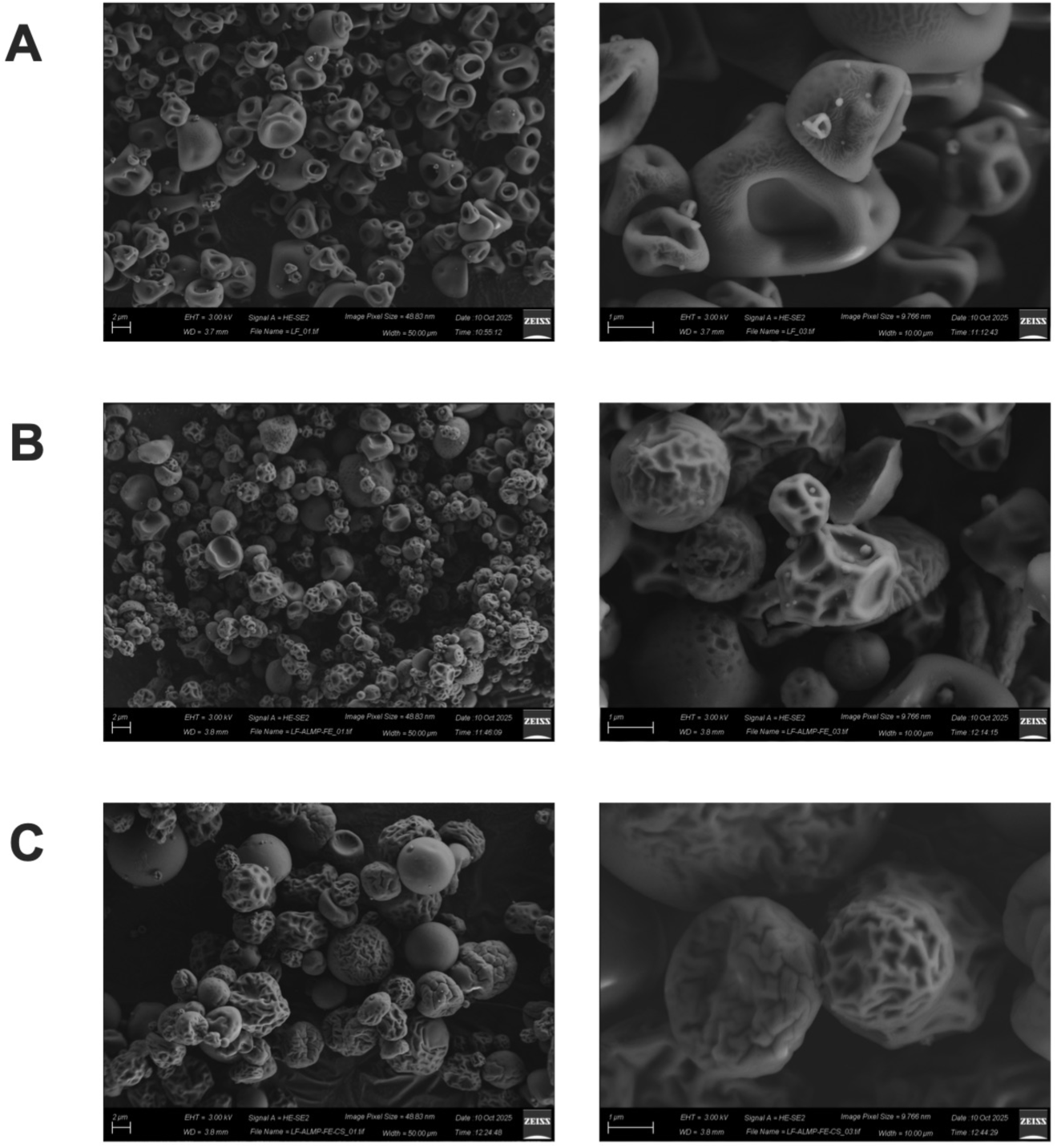
SEM images of spray dried LF (A), LF-aLMP-Fe (B) and CS-LF-aLMP-Fe (C)

After spray drying the LF and total Fe content of the material was determined. The LF loading achieved was 206 mg g^−1^ for LF-aLMP-Fe microparticles, and 154 mg g^−1^ for CS-LF-aLMP-Fe microparticles. The difference in loading is expected and can be accounted for by the addition of the CS material to the spray drying feed solution which effectively lowers the amount of LF that is in solution. The Fe content of the material follows the same pattern at 34 mg g^−1^ for the LF-aLMP-Fe microparticles and 28 mg g^−1^, for the CS-LF-aLMP-Fe microparticles.

### 3.6 In vitro iron bioavailability

Iron absorption takes place in the duodenum in the proximal small intestine (Suzuki et al., 2005). A reverse enteric delivery system could target delivery to the duodenum by preventing release in a neutral pH condition but release in an acid condition, where it would be free to be absorbed by the enterocyte cells that line the small intestine. Iron bioavailability is defined as the soluble fraction at 30 min in 0.1 M HCl (pH 1). This has been shown to correlate with bioavailability models such as CaCO-2 cell uptake (Swain et al., 2003). The solubility fraction of iron in the LF-aLMP-Fe microparticles was 82% (**Fig. 6)**. The solubility fraction of iron in the CS-LF-aLMP-Fe microparticles was similar, at 83%. In acidic conditions the microparticles show burst release, as expected, with nearly all the iron (75-80%) released within the first 15 min.

**Fig 5.**
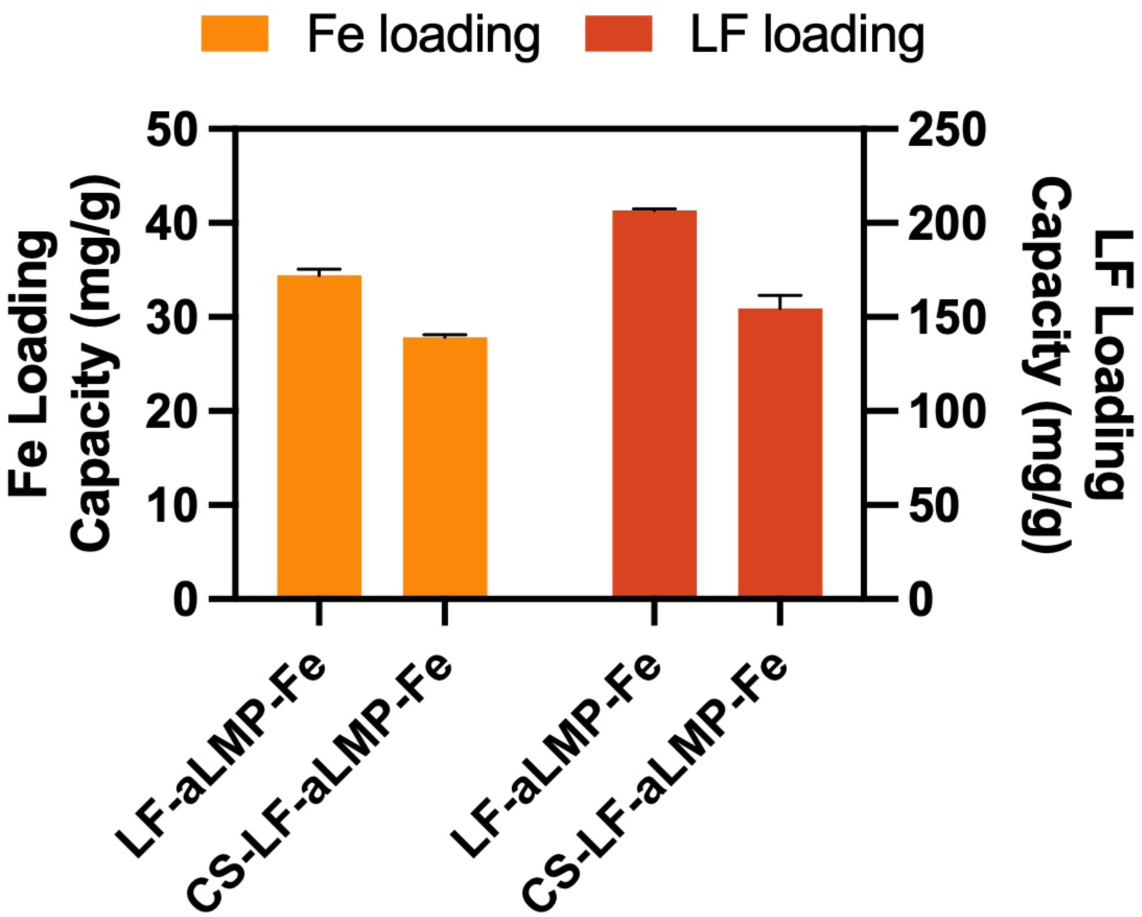
Iron loading (mg g^−1^) and LF loading (mg g^−1^) of the spray dried LF-aLMP-Fe or CS-LF-aLMP-Fe microparticles

**Fig 6.**
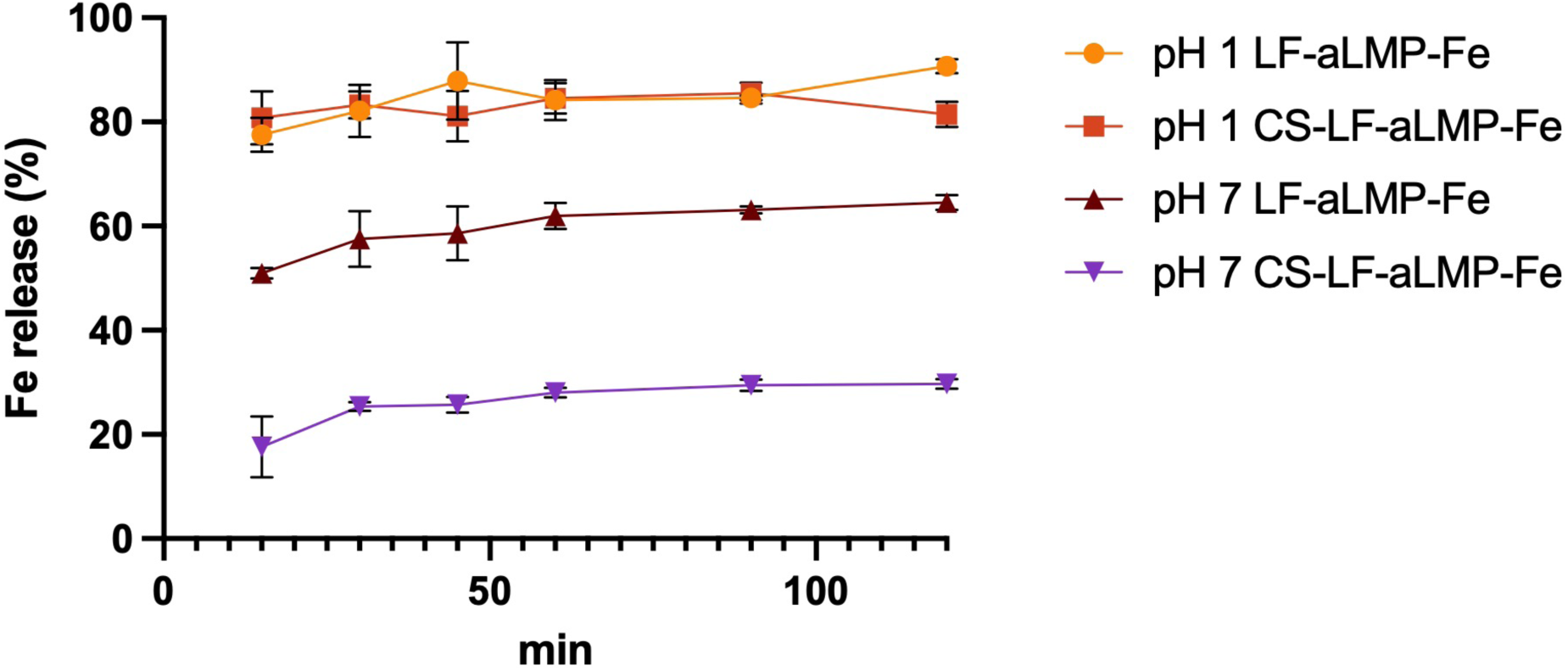
Iron release in pH 1 0.1 N HCl or pH 7 10 mM potassium phosphate buffer from LF-aLMP-Fe or CS-LF-aLMP-Fe microparticles

To reduce metallic undesirable flavors, iron release in the food system or oral condition should be avoided. For both microparticles, the release of iron at pH 7 is much lower, at below 50% at 15 min. Without the addition of CS, the total iron release at pH 7 for the LF-aLMP-Fe microparticles approaches the release in acidic conditions after 120 min, reaching 60%. The CS-LF-aLMP-Fe iron release is slower, starting at 17% in 15 min and plateauing at 30% after 2 h. This is due to the low solubility of CS above pH 4, which prevents the release of the majority of the iron from the core of the microparticle into the solution.

## 4. Conclusion

In this work the co-encapsulation of LF and Fe by aLMP was demonstrated. The LF was trapped in an aLMP-Fe bead through external gelation of the dispersed water phase of a W/O emulsion. The LF-aLMP-Fe microparticles contained 34 mg g^−1^ Fe and 206 mg g^−1^ LF, while the CS-LF-aLMP-Fe microparticles contained 27 mg g^−1^ Fe and 154 mg g^−1^ LF. Encapsulation of the LF decreased proteolysis during simulated gastric and intestinal digestion, demonstrated by intact LF remaining visible in an SDS-PAGE gel. The addition of CS to the LF-aLMP-Fe microparticles reduced Fe release in a neutral pH solution compared to non-CS coated microparticles but did not prevent release in an acidic environment. This encapsulation system shows promise as a method to increase iron absorption compared to FeSO_4_ alone through co-supplementation with LF, while decreasing potential sensory impact on food.

## Supporting Information

Supporting information contains additional figures, discussion and tables that provide additional evidence for the work presented. All raw data associated with the figures can be found at: https://doi.org/10.5281/zenodo.17886326

## CReDIT authorship contribution statement

**Claire Noack:** Conceptualization, Investigation, Data curation, Formal analysis, Writing – original draft, Writing – review & editing. **Peilong Li:** Conceptualization, Methodology, Investigation, Writing – review & editing. **Waritsara Khongkomolsakul:** Conceptualization, Supervision, Funding acquisition, Writing – review & editing. **Yunan Huang:** Investigation, Writing – review & editing. **Alireza Abbaspourrad:**Project Administration, Funding acquisition, Resources, Writing – review & editing, Supervision.

## Generative AI Statement

The authors certify that generative AI was not used in preparing this article. Non-generative AI, such as spelling and grammar checkers in Office 365 and Google Docs, and citation management software, was used. All instances when non-generative AI was used were reviewed by the authors and editors.

## Acknowledgements

This project was funded by the Bill & Melinda Gates Foundation (INV-039533). The authors acknowledge the use of facilities and instrumentation supported by NSF through the Cornell University Materials Research Science and Engineering Center DMR-1719875. CLSM imaging data was acquired through the Cornell Institute of Biotechnology’s BRC Imaging Facility (RRID:SCR 021741), with NYSTEM (C029155) and NIH (S10OD018516) funding. The authors thank Dr. Kelley Donaghy for her valuable assistance in the editing of this manuscript.

