## Supplemental Figures for "Core-shell microparticle encapsulation for pH-responsive and targeted delivery of lactoferrin and ferrous sulfate"

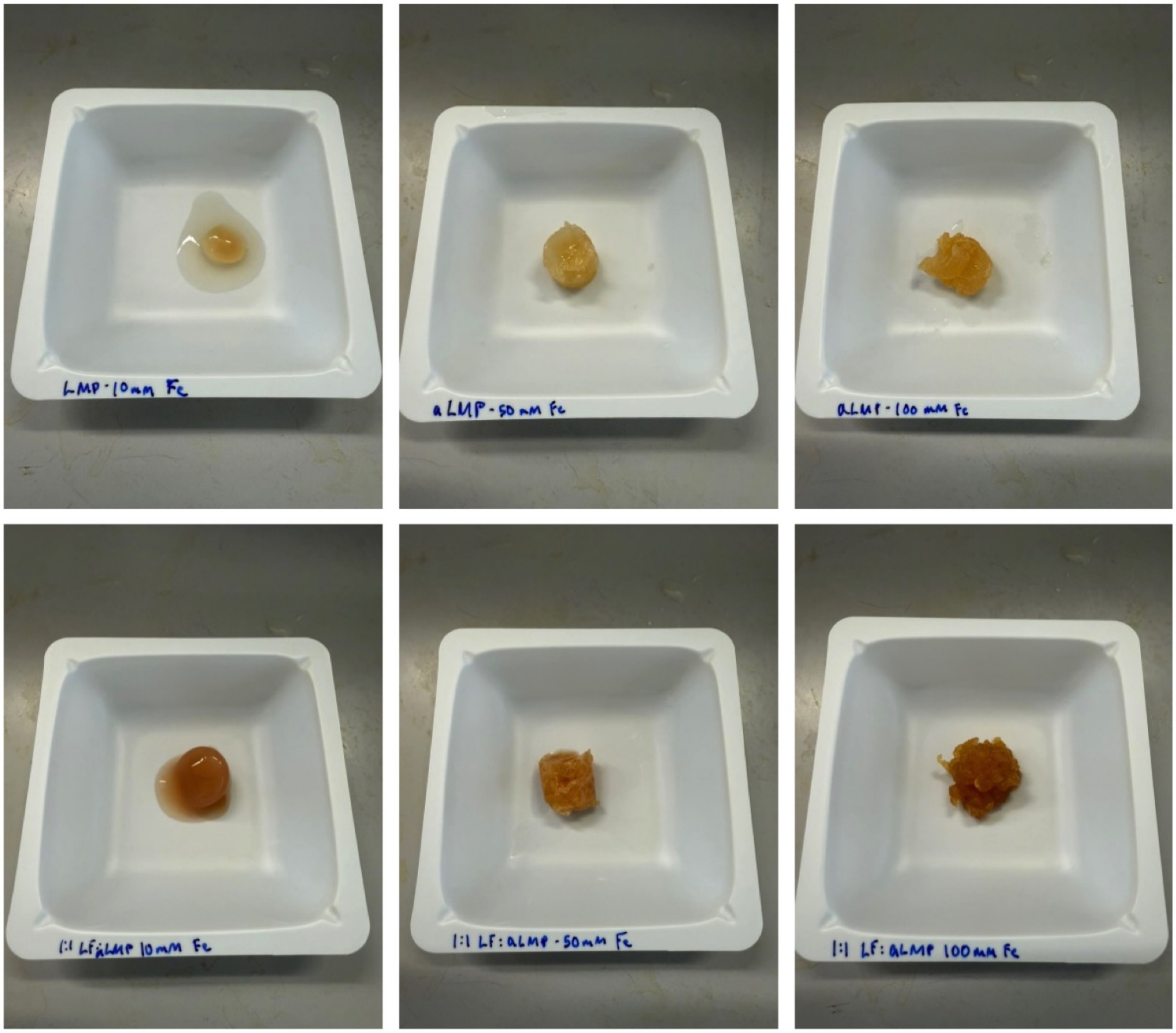


Fig S1. 25 mg $mL^{-1}$ aLMP or 1:1 LF-aLMP-Fe at pH 6 with 10, 50, or 100 mM Fe gel blocks

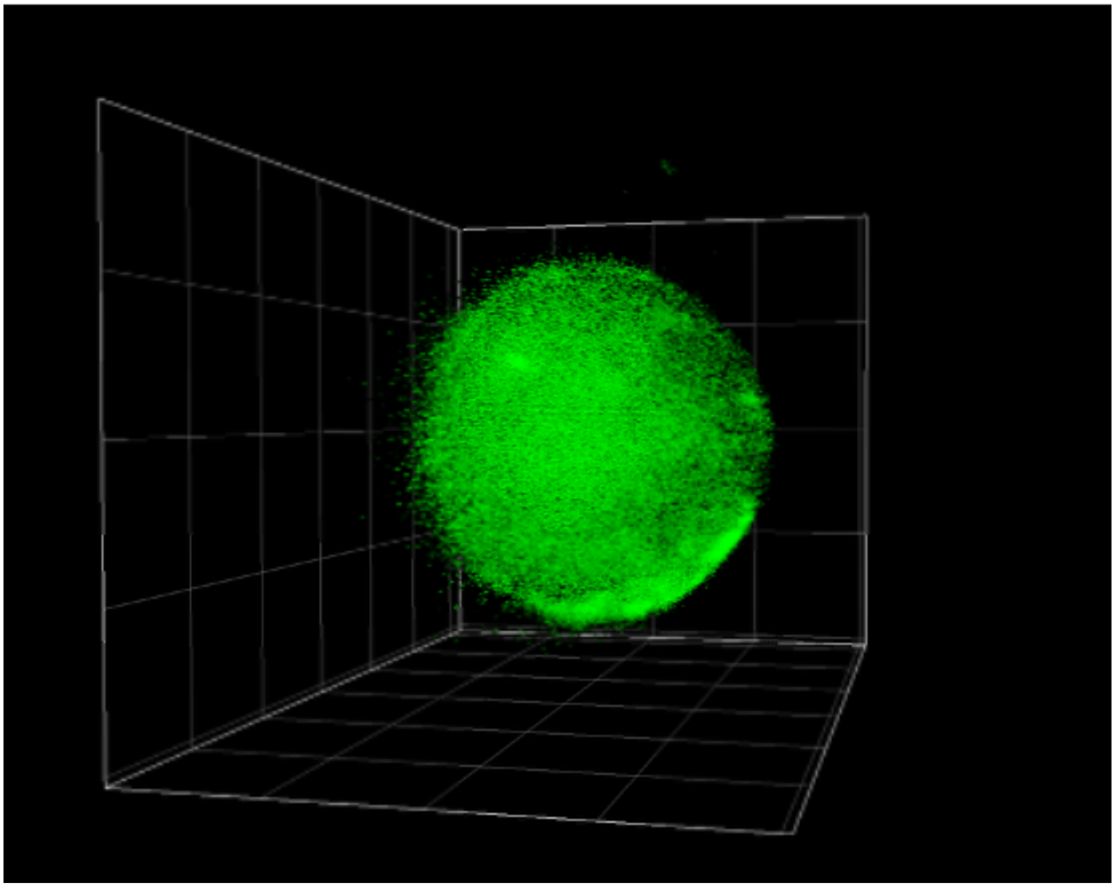

Fig S2. CLSM Z-stack of FITC-labeled LF in water in oil emulsion
